# Generation and Characterization of *Col6a1* knock-in mice: A Promising Pre-Clinical Model for Collagen VI-Related Dystrophies

**DOI:** 10.1101/2025.03.11.642560

**Authors:** Arístides López-Márquez, Carmen Badosa, LLuis Enjuanes, Patricia Hernández-Carabias, Manuel Sánchez-Martín, Bruno Cadot, Zoheir Guesmia, Ioannis Georvasilis, Sol Balsells, Marcos Blanco-Ramos, Emma Puighermanal, Albert Quintana, Mònica Roldán, Valérie Allamand, Cecilia Jiménez-Mallebrera

**Affiliations:** Neuromuscular Unit, Neuropaediatrics Department, Institut de Recerca Sant Joan de Déu, Hospital Sant Joan de Déu, Esplugues de Llobregat, Barcelona, Spain; Center for Biomedical Research on Rare Diseases (CIBERER), Instituto de Salud Carlos III, Madrid, Spain; Department de Genetics, Microbiology and Statistics. University of Barcelona.; Instituto de Investigación Biomédica de Salamanca (IBSAL).; Servicio de Transgénesis. Universidad de Salamanca.; Sorbonne Université-Inserm,Institut de Myologie, Centre de Recherche en Myologie, F-75013 Paris, France; Confocal Microscopy and Cellular Imaging Unit, Department of Genetic and Molecular Medicine, Hospital Sant Joan de Déu, Barcelona, Spain. Institut de Recerca Sant Joan de Déu, Esplugues de Llobregat, Barcelona, Spain; Statistics Department, Institut de Recerca Sant Joan de Déu, Barcelona, Spain; Institut de Neurociències, Universitat Autònoma de Barcelona, Spain; Department of Cell Biology, Physiology and Immunology, Universitat Autònoma de Barcelona, Spain

**Keywords:** Collagen VI-Related Dystrophies, dominant negative mutation, CRISPR/Cas9, mouse model, myopathy, fibrosis, automated image analysis

## Abstract

Collagen VI Related Dystrophies (COL6-RD) are congenital muscle diseases, typically inherited as an autosomal dominant trait. A frequent type of mutation involves glycine substitutions in the triple helical domain of collagen VI alpha chains, exerting a dominant-negative effect on the unaltered protein. Despite this, no prior animal model captured this mutation type. Using CRISPR/Cas9, we generated transgenic mice with the equivalent of the human *COL6A1* c.877 G>A; p. Gly293Arg mutation. We characterized their skeletal muscle phenotype over time, utilizing computer-aided tools applied to standardized parameters of muscle pathology and function. Knock-in mice exhibited early-onset reduced muscle weight, myopathic histology, increased fibrosis, reduced collagen VI expression, muscle weakness, and impaired respiratory function. These features provide adequate outcome measures to assess therapeutic interventions. The different automated image analysis methods deployed here analyze thousands of features simultaneously, enhancing accuracy in describing muscle disease models. Overall, the *Col6a1* Ki Gly292Arg mouse model offers a robust platform to deepen our understanding of COL6-RD and advance its therapeutic landscape.

**Summary Statement:** We generated and characterized over time the first mouse model representing dominant negative glycine substitutions in the alpha chains of collagen VI that are a frequent cause of Collagen VI-Related Dystrophies.

## Introduction

Collagen VI-Related Dystrophies (COL6-RD) encompass a spectrum of ultra-rare myopathies of varying progression and severity from Ullrich Congenital Muscular Dystrophy (UCMD) to Bethlem myopathy (BM) with intermediate phenotypes in between. COL6-RD clinical hallmarks include muscle weakness, distal joint hyperlaxity, proximal joints contractures, and progressive respiratory insufficiency in a proportion of affected children. COL6-RD are orphan diseases lacking an effective therapeutical approach. The only treatment available is symptomatic and based on physical and respiratory therapy, surgery and ventilatory support (Natera de Benito et al., 2020).

Collagen VI is an extracellular matrix protein that is expressed in many tissues including tendon, ligaments and skin which are also affected in individuals with COL6-RD. Collagen VI is synthesized and secreted by tissue specific mesenchymal cells such as fibroadipogenic precursor cells which reside between muscle fibers and are the major source of collagen VI in skeletal muscle (Zou et al., 2008). The three collagen VI polypeptide chains (alpha1, 2 and 3) form a heterotrimer which then associates with another heterotrimer to form dimers and then tetramers which are secreted into the extracellular matrix. In the extracellular space the tetramers associate to form microfibrils with a characteristic beaded-filament aspect which amongst other functions mediates adhesion of the muscle fiber to the surrounding matrix (Cescon et al., 2015; Paco et al., 2015).

COL6-RD is caused by mutations in any of the three major collagen VI genes (*COL6A1*, *COL6A2*, and *COL6A3*). The majority (around 75%) of mutations are *de novo* dominant variants (Allamand et al., 2011; Lamandé et al., 2018). Amongst those, glycine to arginine substitutions in the N-terminus of the triple helical collagen domain are common amongst individuals with the intermediate and milder forms of COL6-RD. The mutant alpha chains carrying these glycine substitutions associate with the alpha chains encoded by the wild type allele and are incorporated into tetramers that will be secreted into the extracellular matrix. The tetramers containing mutant chains interfere with the correct assembly and function of the tetramers formed only by wild-type molecules exerting a dominant negative effect and impairing collagen VI microfibril formation and function. Common missense variants in that region of the triple helix are the ones affecting amino acids 284, 290 and 293 (G284R, G290R and G293R substitutions) in exons 9 or 10 of the *COL6A1* gene. The effect of these glycine substitutions has been characterized at the biochemical level in patient‗s skin fibroblasts, a cell model commonly used for diagnosis and pre-clinical research in COL6-RD (Butterfield et al., 2013; Jimenez-Mallebrera et al., 2006).

We and other groups have reported promising *in vitro* results with either siRNAs, ASOs or CRISPR/Cas9 (Noguchi et al., 2014; López-Márquez et al., 2022; Brull et al., 2024);) aimed at reducing the expression of the G284R and G293R mutated alleles and thus their detrimental effect. These approaches demonstrated significant silencing of the mutant allele that was accompanied by an increase in collagen VI matrix deposition, an improvement in the architecture of collagen VI microfibrillar network and the recovery of various subcellular alterations (Castroflorio et al., 2022).

Research in collagen VI deficient primary cells and in the existing animal models for recessive and dominant COL6-RD has identified various underlying molecular alterations including fibrosis and extracellular matrix remodeling, impaired autophagy, mitochondrial dysfunction, muscle cell atrophy and intrinsic stem cell defects. Several of those studies were conducted in fibroblasts and/or muscle biopsies from COL6-RD affected individuals carrying one of those frequent glycine substitutions. However, there is no reported mouse model that represents that typical type of mutation. Using CRISPR/Cas9 we have generated a knock-in mouse model that harbors the C*ol6a1* c.874 G>A; p. G292R mutation that mimics the human mutation *COL6A1* c.877 G>A; p. G293R. We refer to these mice as C*ol6a1* Ki G292R. Our purpose is to use this model to investigate mutation-specific and general disease drivers and modifiers, derive useful read-outs to guide future pre-clinical studies and validate potential therapies. Here we characterize systematically this novel mouse model deploying different qualitative and quantitative tools on standardized muscle pathology parameters. *Col6a1* Ki G292R mice show hallmarks of COL6-RD such as fibrosis, partial collagen VI deficiency and reduced limb strength, provide some new insights into the disease and overall constitute a valuable model for COL6-RD.

## Results

### Generation of C*ol6a*1 knock-in mice using CRISPR/Cas9

We designed a gRNA targeting exon 10 using Breaking-Cas tool (Oliveroset al. 2016) and a 200bp single stranded DNA template (ssODN) that contained the desired muted base (c.874 G>A), (Fig 1A). These were microinjected together with SpCas9 into C57BL6/J zygotes and implanted in pseudopregnant females. Edited founders were identified by PCR amplification (Supp. Table 1) and those carrying the desired alleles, were crossed five generations with wild-type C57BL6/6J to eliminate possible unwanted off-targets. Heterozygous mice were re-sequenced and crossed with C57BL6/6J wild-type animals to generate the BL/6J-Col6A1em1(c.874G>A) ^Sal^ mouse colony which we refer to as *Col6a1* Ki G292R.

**Fig 1.**
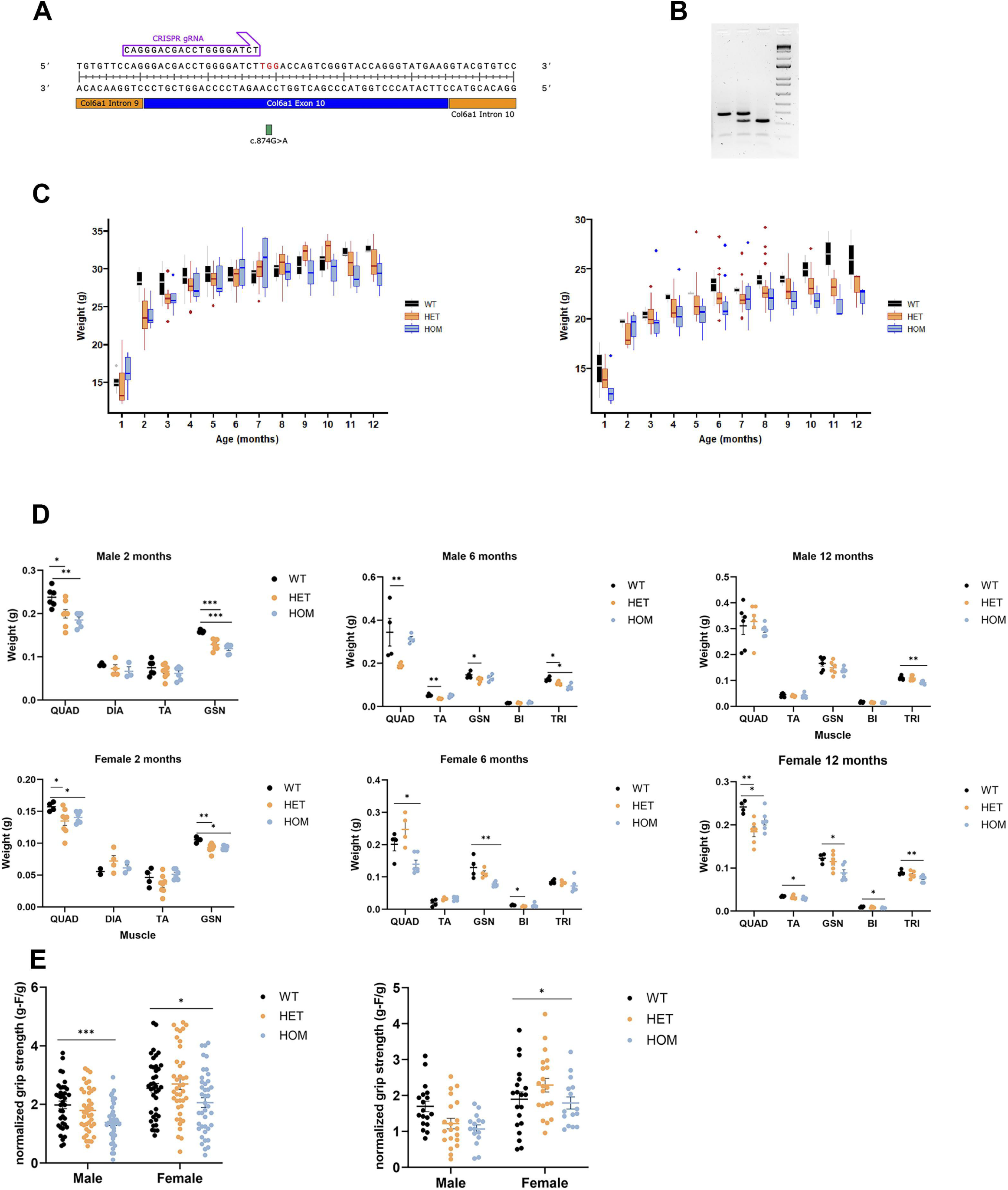
A. Position of gRNA used to generate the Col6a1 Ki mice relative to the position of the introduced mutation in the col6a1 gene. B. Representative agarose gel electrophoresis of the digestion fragments corresponding to the wild-type, heterozygous and homozygous genotypes. C. Graphs representing the progression of the body weights for male and female mice over a 12-month period (a linear regression model was used, see materials and methods, median and interquartile ranges are represented). D. Weights of different muscles from male and female mice are shown separately at 2, 6 and 12 months of age. E. Grip strength of the forelimbs was assessed at 6 and 12 months in male and female separately. In D and E, means and SEM are represented. One-way ANOVA was used followed by Tukey’s HSD as a post-hoc test to determine statistical significance; between 4 and 8 mice were assessed of each sex and age. *= p< 0.05, ** = p < 0.1; *** = p < 0.001, **** = p< 0.0001.

### *Col6a1* Ki G292R mice have retarded growth, muscle atrophy and weakness

Mutant mice moved and bred normally and had a normal life span. They were not overtly different from wild-type mice. We obtained the expected numbers of each genotype in the offsprings according to Mendelian inheritance and a comparable number of animals of each sex. Regarding the growth of the mice in terms of body weight, we observed differences between genotypes (and sexes) that we explored using piecewise and linear regression models for males and females separately between the ages of 1 month and 12 months. We observed that with the available data the progression of the body weights could be divided into an initial phase of faster growth, followed by a second phase of slower growth. We applied a linear regression model with segmented relationship that estimated that the breakpoint between the two phases occurs at around 2.2 months (95% CI 2.01-2.47). Then, we analyzed each period separately and studied how age, sex and genotype explained weight (Fig 1C). The values of the resulting coefficients and adjusted p values are included in Suppl. Table 2. In male mice, the wild-type group grew more rapidly in the first phase than heterozygous and homozygous mice (adjusted p-value < 0.05 in both cases represented by the coefficient Age*Genotype HET or HOM in Suppl. Table 2), indicating that they reached their adult weight faster. In the second phase we also obtained significant results indicating that heterozygous mice start from a lower baseline (corrected p-value < 0.05, represented by the coefficient Age*Genotype HET in Suppl. Table 2), but all mice eventually reach similar weight by 12 months. In female mice, the analysis did not support differences in growth rate between genotypes (Suppl. Table 2).

We analyzed the weight and morphology by hematoxilin and eosin staining (H&E)) of the liver, kidney, brain, lung, intestine and spleen of 6- and 12-months mice separately in males and females to reveal any potential subjacent non-skeletal pathology but no major changes in the normalized weight of the organs or tissue morphology related to the genetic modification were observed (data not shown).

Regarding the weight of individual muscles, *col6a1* Ki muscles weighed less than the muscles of their wild type littermates and remained lighter up to 12 months (Fig 1D).

We tested the strength of the forelimbs of the individuals to evaluate whether the presence of the pathogenic variant also affects the ability to grip and generate strength. Despite the variability in the measurements between individuals, we determined a significant reduction in strength between wild-type and homozygous mice at 6 and 12 months which was more pronounced in males than females (Fig 1E). In heterozygous mice the measurements were on average lower than in the wild-type group, but the differences were not statistically significant. Additional functional assays assessing endurance or coordination, such as the Rotarod test, exhibited high variability and did not reveal any significant genotype-related differences (data not shown).

### Respiratory function evolves differently in *Col6a1* Ki G292R mice compared to wild-type mice

We investigated respiratory function in *Col6a1* Ki G292R mice using whole-body plethysmography (WBP), a non-invasive technique that allows recording several indicators of respiratory function. We recorded data for male and female wild-type, heterozygous and homozygous mice separately at two points in time, 6 and 12 months. We applied a linear mixed model to investigate the effect of three factors: genotype, sex and moment in time, and their interactions. The values of the resulting coefficients and adjusted p values for each parameter are listed in Suppl. Table 2. Some parameters (End Inspiratory Pause, EIP, and End Expiratory Pause, EEP) were excluded because of the presence of outliers (data not shown). Data for the other parameters are summarized in Fig 2.

**Fig 2.**
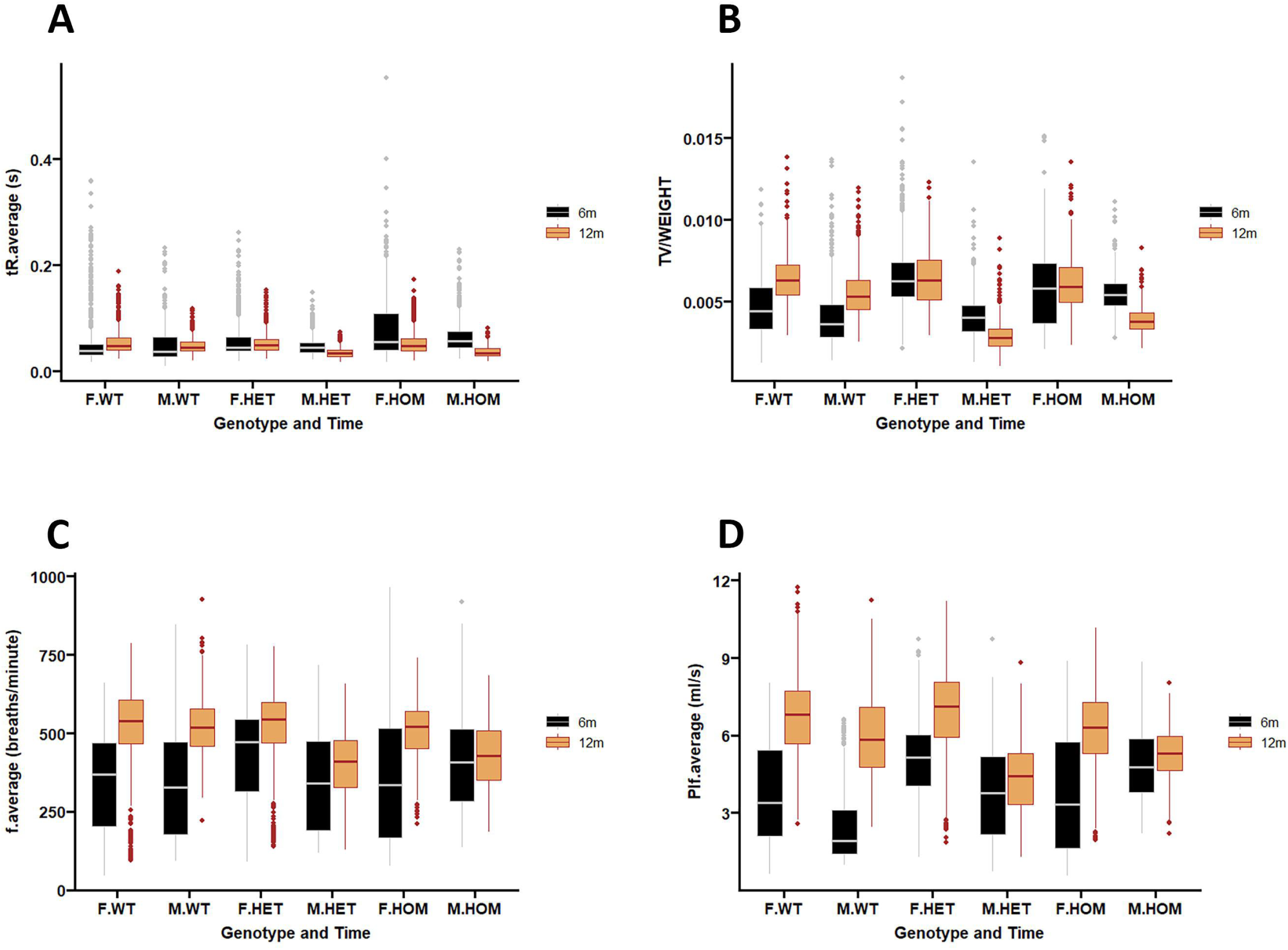
Distribution of respiratory function parameters assessed by whole body plethysmography analyzed by sex and genotype at 6 and 12 months (see Materials and Methods for details of the statistical analysis and Suppl Table 3 for adjusted p-values of the calculated coefficients). TR: relaxation time, TV/Weight: Tidal volume/weight of the animal, f: respiratory rate and Pif: Peak Inspiratory Flow.

At 6 months, the average relaxation time (RT) was significantly longer (p < 0.001) in homozygous mice relative to wild-type mice. When we analyzed the changes in the different respiratory parameters between 6 and 12 months, we observed that in wild-type mice (regardless of the sex) there was a significant increase (p<0.001) in tidal volume (TV/weight of mice), respiratory rate (average F) and Peak inspiratory flow (Pif). However, this increase was not confirmed in heterozygous or homozygous mice. The average pause index (Penh) decreased (p<0.001) in wild-type mice while in heterozygous or homozygous male mice this change was positive (p<0.001). In summary, WBP showed that respiratory function evolves differently in *Col6a1* Ki mice in comparison to their wild-type littermates and this difference is more marked in males than females.

### *Col6a1* Ki G292R mice show early myopathic changes including increased fibrosis in various muscles limb muscles and the diaphragm

We evaluated qualitatively the skeletal muscle overall pathology using H&E in different muscles (quadriceps, tibialis anterior, gastrocnemius, soleus, biceps, triceps and diaphragm), across the three genotypes at different ages (2, 6 and 12 months) in male and female mice. First, two examiners conducted blindly a qualitative assessment of the histological preparations and then one of them, with more experience in muscle pathology, analyzed them in more detail. We observed mild pathological changes at 2 months in heterozygous and homozygous mice in the different limb muscles examined and diaphragm, consisting of increased presence of internal nuclei, variation in fiber size towards smaller fibers and roundness of muscle fibers. At 6 months of age, we observed a similar degree of pathology in all muscles including triceps and biceps. In some cases, the pathology appeared more marked in the homozygous than in the heterozygous mice, but this was not always the case and depended on the individual mouse and the muscle (Fig 3). At 12 months the pathological changes persisted but did not seem to worsen with age (Supp. Fig 1). In the diaphragm the differences between heterozygous and homozygous and wild type mice were more apparent than in younger mice although the pathology of the diaphragm is difficult to interpret because of the technical difficulty in orienting the muscle fibers and in avoiding artifacts. In summary, this qualitative morphological analysis revealed mild pathological changes from 2 months of age in all limb muscles and in the diaphragm that were compatible with a myopathic process.

**Fig 3.**
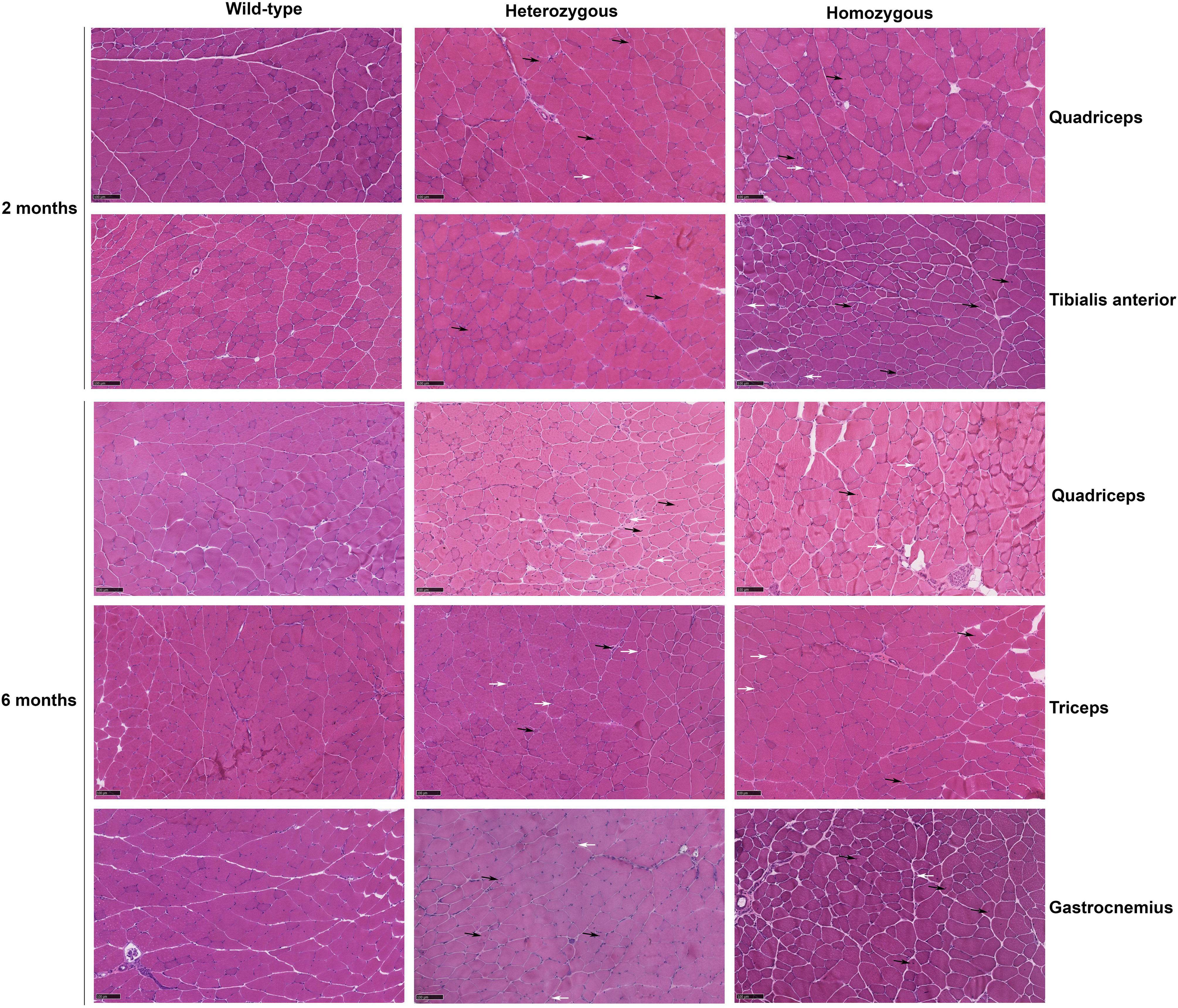
Frozen sections of different muscles were stained with hematoxilin and eosin (H&E). Myopathic features include fibre size variability with small atrophic fibres (white arrows) and increased internal nuclei (black arrows). Scale bar= 100µm.

To quantitate and statistically evaluate muscle pathology we applied an automated purpose-built pipeline on whole section reconstructions of the different muscle groups and diaphragm from 6 months old mice (see M&M) analyzed separately by sex. This allowed us to obtain many fibres for each muscle (ranging from 893 to over 9,000 features). We focused on two parameters: the minimum Feret‗s diameter is a robust measurement of muscle fiber size. It avoids errors due to the orientation of the sectioning angle and is widely used to describe the variation in muscle fiber diameter in animal models of muscular dystrophy. On the other hand, the presence and number of internal nuclei, an indicator of muscle degeneration and regeneration, is also frequently used in this context.

Regarding the Feret‗s diameter, although we did observe in some cases significant differences in some muscles after adjusting the p-values for multiple test comparison (data not shown), there was not a global consistent pattern across muscles and genotype. For that reason, we looked at the coefficient of variation and its confidence interval (95% CI). Using this as an indicator of fiber size variability we obtained some differences (we considered non-overlapping CI to be relevant) but only in some cases: female gastrocnemius and female tibialis anterior and male triceps (SupplTable 3).

For the number of internal nuclei, we looked at the distribution of the % of fibres with 0 or more internal nuclei. There were significant and relevant differences between genotypes only in male tibialis anterior (adjusted p value < 0.001, effect size r > 0.1). In this case, in heterozygous mice there was a significant and relevant lower proportion of fibres with 0 internal nuclei (87. 3%) than in wild-type mice (96.1%).

These results indicate that when a sufficiently large number of fibres are analysed, there are no objective and quantitative differences in the diameter of the fibers or the number of internal nuclei between genotypes that are consistent across different muscles even if these features are present.

To determine the presence and extent of endomysial fibrosis we stained sections using the Picro Sirius Red stain (PS) and measured the percentage area occupied by collagen fibrils. We analyzed quantitatively the diaphragm, quadriceps and tibialis anterior in 2-, 6- and 12- months mice by applying an automatic image analysis pipeline that we programmed with FIJI- ImageJ Software. A significant increase in collagen deposition was observed from 2 months in the three muscles in heterozygous and homozygous mice relative to wild type littermates (Fig 4). The same trend was observed at 6 and 12 months including the triceps, which were not collected at 2 months. In some cases, there were significant differences as well between heterozygous and homozygous mice (for example in diaphragm at 2, 6 and 12 months, Fig 4). In summary, there was pathological fibrosis in all muscle tested in Knock-in mice.

**Fig 4.**
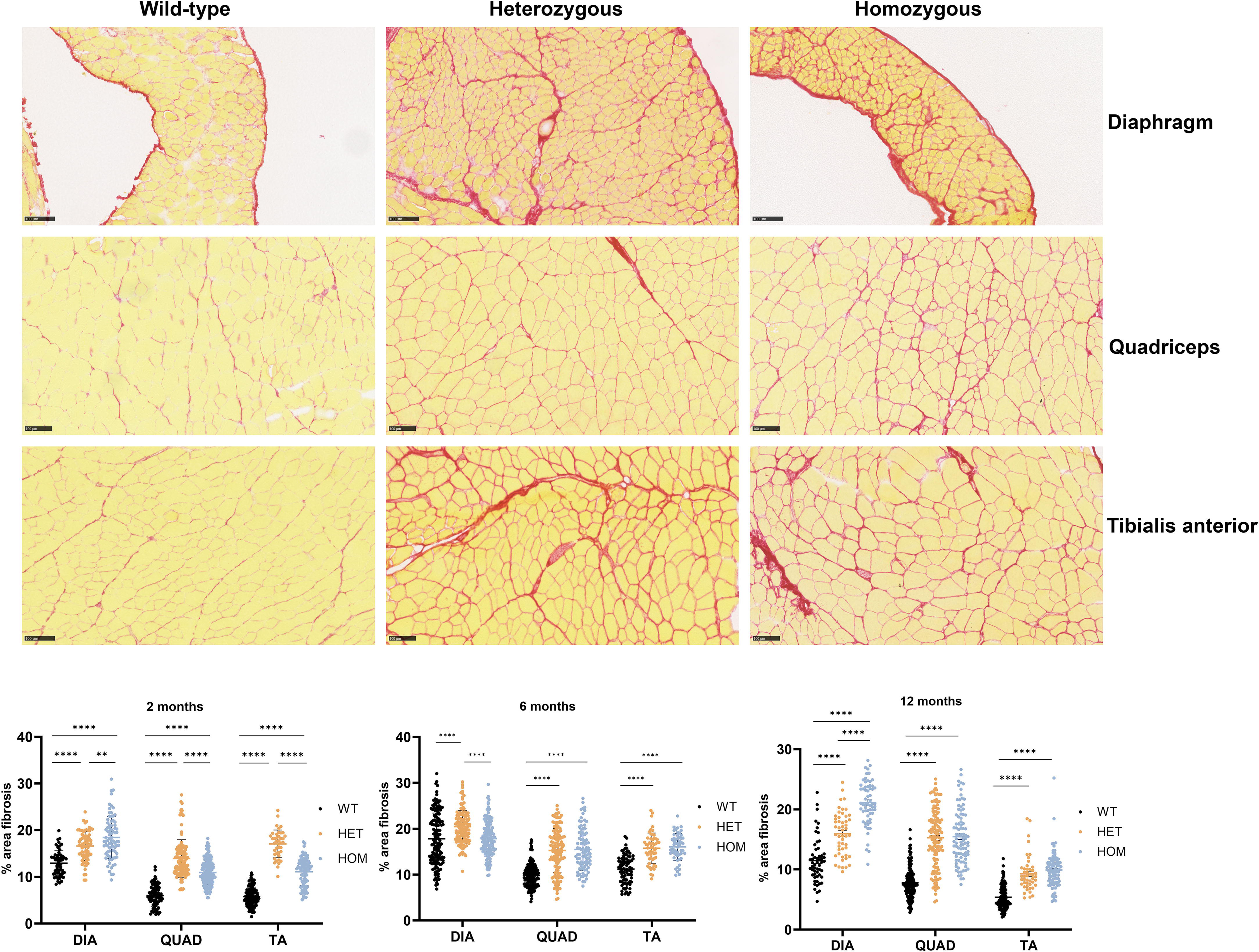
Picro Sirius Red (PS) staining of frozen sections was used to reveal collagen deposition as an indicator of fibrosis. Scale bar= 100µm. The % area of fibrosis were quantified and are represented in violin plots for mice aged 2, 6 and 12 months. *= p< 0.05, ** = p < 0.1; *** = p < 0.001, **** = p< 0.0001.

### Mouse diaphragm shows the highest expression of *Col6a1* and the expression of *Col6a2*, *Col6a3*, *Col6a5* and *Col6a6* is unchanged in *Col6a1* Ki mice

We analyzed the expression of *Col6a* transcripts by digital droplet PCR in different muscles in 2 and 12-month-old mice. On one hand, we measured total *Col6a1*, *Col6a2*, *Col6a3*, *Col6a4* and *Col6a5* mRNAs levels and on the other hand we used allele specific probes to determine the expression of the *Col6a1* wild-type and mutated alleles and investigate if there was any deviation from the expected 50% expression of each allele. We did not observe significant differences between males and females in total *Col6a1* expression levels, so data were pooled. In some muscles, the levels of *Col6a1* transcript were higher in heterozygous and/or homozygous mice than in wild type mice although in most cases those differences were not significant (Fig 5). When we compared the copies/ul of *Col6a1* across the different muscles tested in wild type mice, we found that diaphragm was the tissue which expressed the highest levels followed by soleus, tibialis anterior, quadriceps and Gastrocnemius (Fig 5). The same results were observed at 12 months (data not shown).

**Fig 5.**
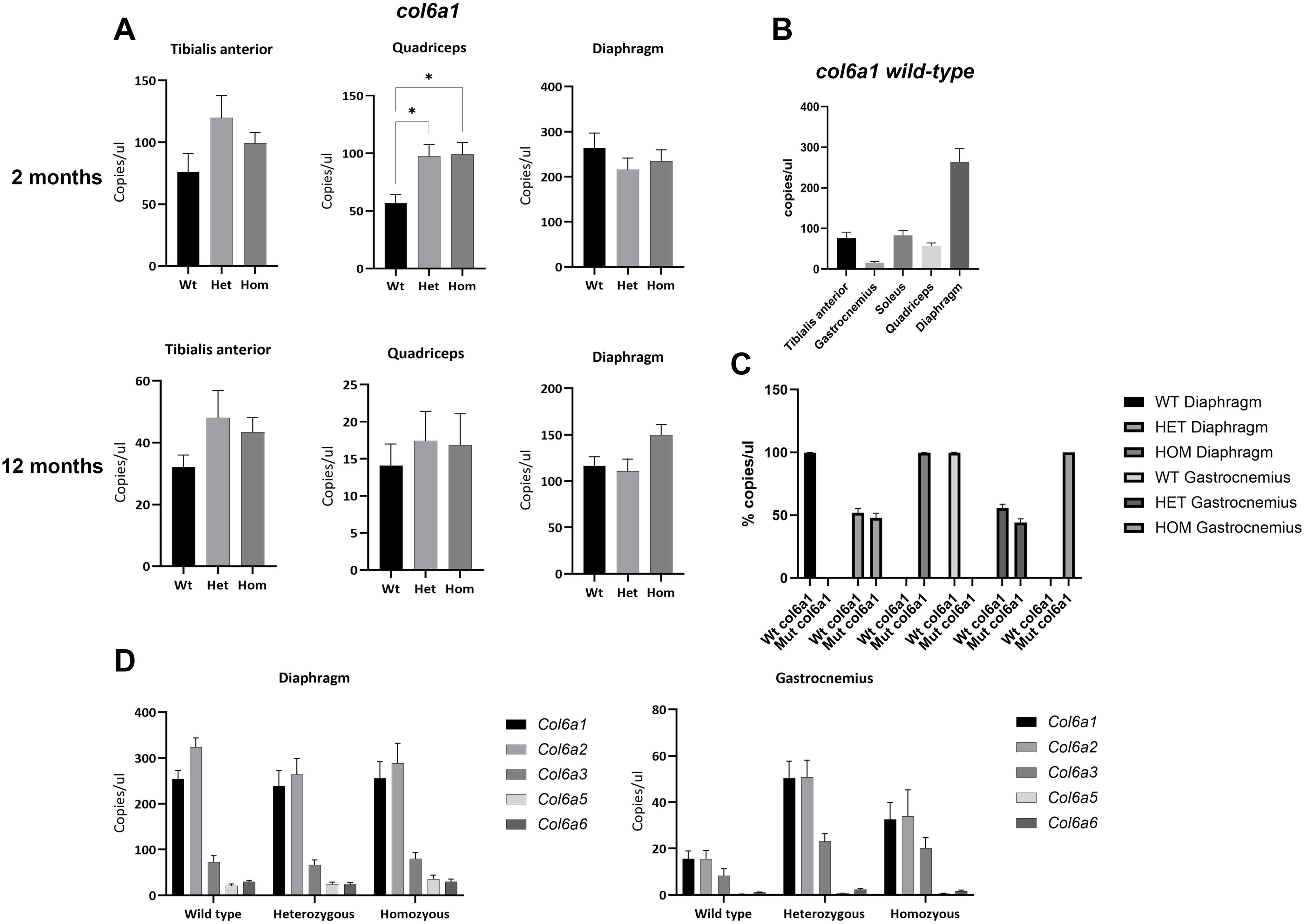
Expression of Col6a transcripts: A. Col6a1 expression in tibialis anterior, quadriceps and diaphragm of wild-type, heterozygous and homozygous 2 (Top) and 12 (Bottom) months-old mice (n =5 wt, 6 het, 6 hom). B. Comparison of total col6a1 levels in different muscles in wild-type mice at 2 months (n=5). C. Abundance (copies/µl) of col6a1 wild-type (Wt) and mutant (Mut) col6a1 transcripts in diaphragm and gastrocnemius in 2 months old wild-type (wt, n=5), heterozygous (het, n=6) or homozygous (hom, n=6) mice D. Expression of col6a1, col6a2, col6a3, col6a5 and col6a6 in diaphragm and gastrocnemius. Two-way ANOVA was performed to compare the different groups, followed by Tuke’s multiple comparison test for two –way comparisons between genotypes. Data from two experiments are represented as mean ± SEM. *= p <0.05, **= p<0.01 *** = p <0.0005; ****= p <0.0001.

Expression of *Col6a5* and *Col6a6* has been described in a variety of mouse tissues (Fitzgerald et al., 2008). We assessed the expression of *Col6a2*, *Col6a3*, *Col6a5* and *Col6a5* in two representative muscles (Gastrocnemius and Diaphragm) in 2 months-old mice. Expression of the transcripts for *Col6a1* and *Col6a2* was much higher than the *Col6a3*, *Col6a5* and *Col6a6* transcripts in both muscles tested but no differences were detected between wild type and knock-in mice (Fig 5).

Using allele-specific probes we determined the ratio of wild type (a G at position 872) vs mutant (an A at position 872) *Col6a1* allele. The percentage of each allele in heterozygous mice only slightly deviated from the expected 50% ratio (Fig 5).

### Collagen VI is reduced in *Col6a1* Ki muscles and diaphragm at different ages

To detect and semi-quantitate collagen VI alpha chains, we analyzed by SDS-PAGE and Western blot the presence and relatively abundance of the alpha1 (VI) polypeptide in total protein extracts of different muscles at 2, 6 and 12 months. Firstly, we compared males and females but found no significant differences. Thereafter, data from male and female mice were analyzed together. We observed a significant reduction in the relative levels of alpha1 (VI) in all the muscles tested (triceps, diaphragm, gastrocnemius and tibialis anterior) at least at one of the three ages analyzed (Fig 6).

**Fig 6.**
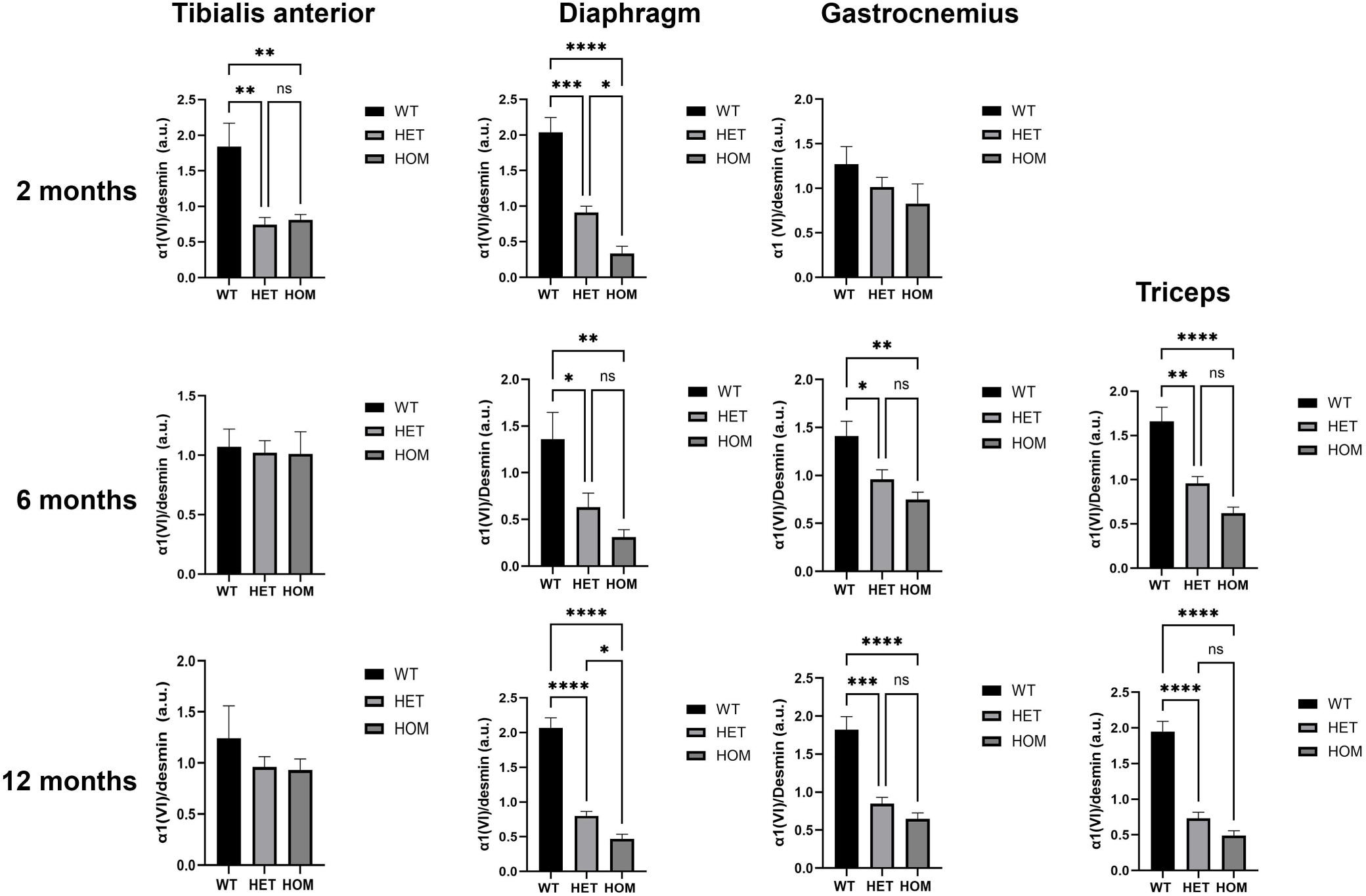
Bars graphs showing the relative abundance of a1 (VI) in triceps, diaphragm, gastronecmius and tibialis anterior muscles of wild-type, heterozygous and homozygous mice at different ages (between 3 and 6 mice per genotype at each stage). Col6a1 was detected with a polyclonal antibody against Collagen Type VI (70R-CR009x). Data were analysed using a One-way ANOVA followed by Tukeýs multiple comparison test. Data are mean ± SEM *p=0.01; **p<0.001; ****p<0.0001.

Muscle cryosections from 12 months old mice were doubly-immunostained for collagen VI and perlecan, since this is the protein that we use as control of basal lamina integrity in human biopsies. A minimum of two slides (4 sections) per muscle and mouse were processed from two separate positions of the muscle block. Confocal microscopy whole section mosaics were captured, muscle fibers segmented and the fluorescence intensity around each muscle fiber was automatically measured. The automation allowed us to quantify a very large number of muscle fibres per muscle. An initial analysis of collagen VI intensity separated by sex in some muscles did not detect significant differences and therefore the analysis was performed irrespectively of sex. We compared for each the collagen VI/perlecan ratio between genotypes. In all muscles analyzed (biceps, diaphragm, quadriceps, gastrocnemius, soleus and triceps), except tibialis anterior, the collagen VI/perlecan ratio was significantly decreased in heterozygotes and homozygous mice relative to the wild type littermates (Fig 7). The greatest reduction in terms of mean fluorescence intensity and the extent of the statistical significance were observed in diaphragm, quadriceps and soleus (Supp. Fig 2 and Supp Table 4).

**Figure 7.**
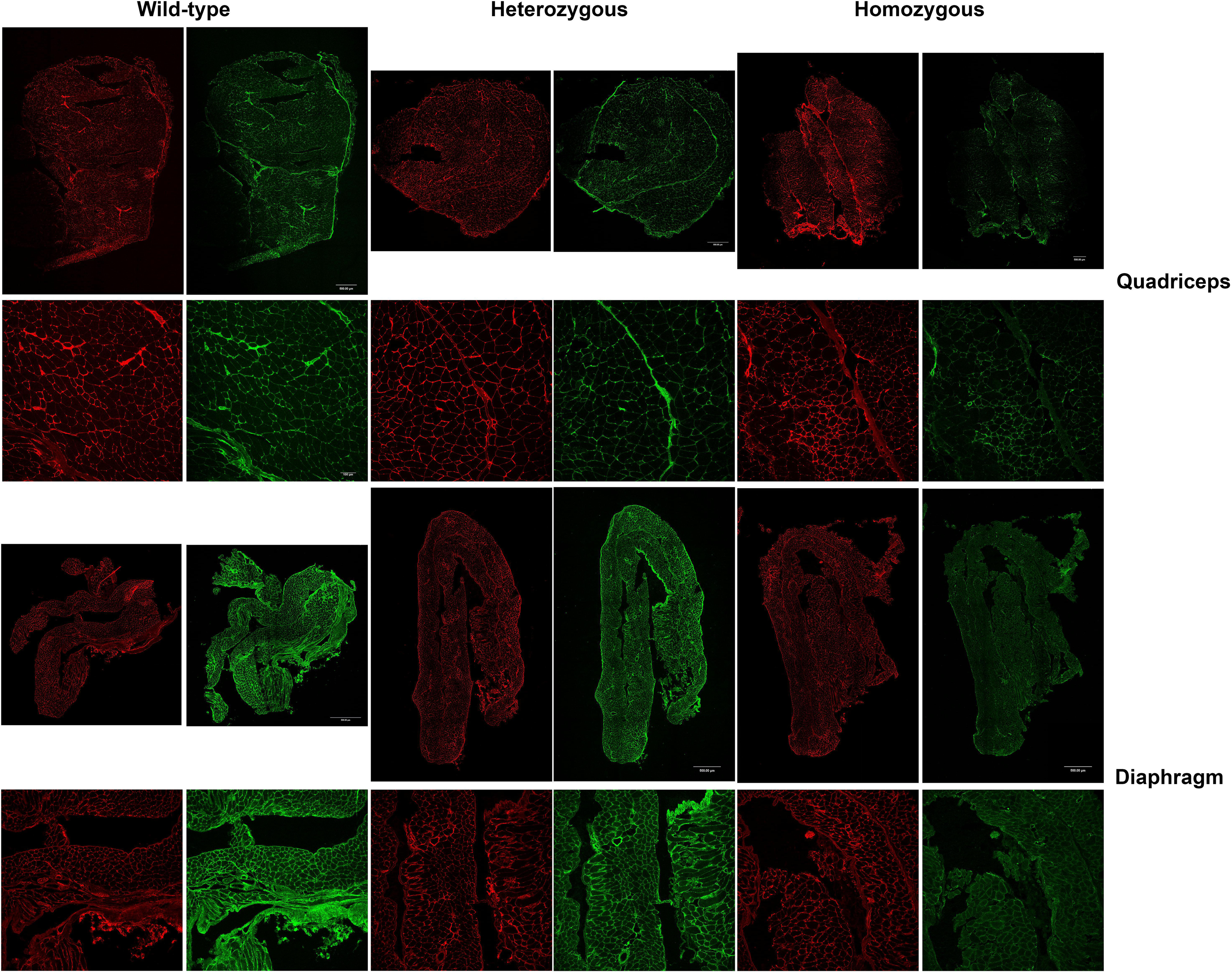
Collagen VI (green) and perlecan (red) double immunofluorescence and confocal microscopy of 12 months old muscle sections from quadriceps and diaphragm. Top row for each muscle represents the whole mosaics (scale bar 500µm) and the bottom row a representative individual field (scale bar 100µm).

## Discussion

In this work we describe a new mouse model of Collagen VI related dystrophies (COL6-RD), in which CRISPR-Cas9 technology was used to introduce a missense mutation in exon 10 of the C*ol6a1* gene. This mutation affects a glycine residue (aminoacid 292) in the N-terminus of the Col6 alpha1 chain that is the equivalent to amino acid 293 in the human protein. The Glycine 293 to Arginine transition as well as other equivalent transitions in that region of the alpha 1 chain are a common type of mutation in affected individuals and therefore this novel mouse model is relevant for COL6-RD. To our knowledge, this is the first knock-in mouse model of COL6-RD with this type of mutation that is associated with the intermediate and milder forms of COL6-RD.

We provide evidence of pathological and functional alterations in mutant mice compatible with a myopathic process, including reduced muscle mass, excess deposition of endomysial collagens, as an indication of fibrosis, from an early age, a partial deficit in collagen VI protein levels and localization at the basal lamina, reduced strength and impaired respiratory function. These features are also observed in individuals with COL6_RD and the *COL6A1* Gly293Arg mutation (Paco et al., 2012, Paco et al., 2013; Butterfield et al.,2013).

Other mouse models for dominant negative mutations include mice with deletion of exon 16 of *Col6a3* (Pan et al., 2014), mice with deletion of exon 5 of *Col6a2* (de Greef et al., 2016) and humanized mice carrying the variant *COL6A1* 930+189C>T (Bolduc et al., 2024). In general, these models show similar features to the *Col6a1* Knock-in mice reported here in terms of reduced muscle mass and limb strength and mild myopathic features including interstitial fibrosis (Pan et al., 2014).

Pulmonary function is a relevant disease outcome measure in the context of COL6-RD. In affected individuals, this is regularly assessed with the Force Vital Capacity (FVC) measurements. In individuals with the intermediate and mild forms of COL6-RD, associated with the *COL6A1* het. c. 877 G>A mutation, the predicted FVC is on average around 60% and 75% of the predicted value and it was estimated that by the age of 20 years old, 30% of individuals classified as intermediate UCMD were using nocturnal ventilation in the form of BiPAP (Natera de Benito et al., 2020). Whole body plethysmography is a very valuable experimental approximation to evaluate respiratory function in animal models. In the other mouse models of dominant collagen VI mutations analysis of pulmonary function was not reported, but it has been reported in models of other congenital muscular dystrophies. For example, in laminin-2 deficient *dy^2J^* mice treated with an anti-apoptotic drug respiratory rate measured by whole body plethysmography was a sensitive monitoring outcome measure (Yu et al., 2013). Similarly, we believe that the changes we found over time in *Col6a1* G292R knock in mice in various respiratory parameters represent very valuable pre-clinical physiological readouts to evaluate the effect of therapeutic interventions in this mouse model.

When we analyzed the expression levels of different *Col6a* mRNAs we observed in some muscles an average increased expression of *Col6a1* transcript in knock-in mice. We also observe this increase when we analyze the RNA from fibroblasts from individuals carrying the equivalent *COL6A1* variant (personal observation). An over-deposition of *Col6a5* and *Col6a6* in the extracellular matrix by immunofluorescence was described in the *Col6a1* exon 16 deleted mouse model (Pan et al., 2014). However, we did not detect changes in these two *Col6a* chains at the transcriptional level, which suggests that the over-expression of *Col6a5* and *Col6a6* may be regulated at the post-translational level.

Using immunoblot we evidenced a significant and consistent reduction in the relative levels of the Col6a1 chain in different muscles in knock-in mice. Differences in the results obtained at the transcript level may be explained by the reduced stability or solubility of the protein chain that is not reflected in transcript levels. Although immunoblot is not a quantitative technique it is nonetheless a useful tool to assess changes in collagen VI. The partial reduction in collagen VI intensity around muscle fibers by immunofluorescence and confocal microscopy is also a relevant finding because it is comparable to what it is usually found in COL6-RD affected individuals with dominant mutations including the *COL6A1* c.877 G>A mutation.

Although in humans the *COL6A1* c. 877 G>A mutation is not present in homozygosity here we analyzed in parallel the heterozygous and the homozygous *Col6a1* G292R genotypes. All the alterations were present in the heterozygous state although more aggravated in homozygous mice (for example the decrease in grip strength).

One of the main contributions of this study is that we strived to develop novel or improved quantitative image analysis pipelines to acquire and extract information from large datasets of morphological or intensity measurements that would represent more faithfully the pathology of this mouse model and that can be applied to other muscle disease models. In the past, we have also applied quantitative image analysis to study other muscle proteins such as dystrophin (Codina et al., 2023). We found that muscle fiber segmentation is a key step in this process, which can be automated using existing tools such as Cellpose3.0, but that can be improved by manual curation. Interestingly, we found that some morphological alterations such as the increase in internal myonuclei that were apparent under the microscope when examining a relatively small number of muscles and optical fields were not confirmed by the statistical analysis of hundreds or thousands of muscle fibres after size effect correction. COL6-RD is not characterized by muscle fiber degeneration or regeneration so perhaps it is not surprising that internal nuclei was not confirmed as a differential parameter between genotypes. In contrast, we and others have described at the microscopic and ultrastructural level the presence of atrophic fibers in COL6-RD human muscle biopsies (Paco et al., 2012). There are many possible explanations for this apparent discrepancy between observational and quantitative results including the variability between individual mice and the effect of the observer. The observed morphological alterations could nonetheless be biologically relevant, although subtle. In any case, it highlights the importance of conducting extensive and quantitative analysis in disease models particularly when they are used as a platform for therapeutic pre-clinical research.

One of the most robust pathological findings was the fibrosis observed from an early age in heterozygous and homozygous *Col6a1* Ki mice in all muscles analyzed. Fibrosis has also been described in other COL6-RD mouse models and in other murine models of Congenital Muscular Dystrophy (Pan et al., 2014; Mohassel et al., 2023). It is also a frequent finding in muscle biopsies from individuals with COL6-RD. Extracellular matrix remodeling and fibrosis is also a commo finding of various transcriptomic studies of collagen VI deficient muscle and fibroblasts (Paco et al., 2013). Recent studies have shown dysregulation of the TGFβ pathway leading to fibrosis and impaired regeneration after muscle injury in *Col6a2* knock-out mice as an early disease driver (Mohassel et al., 2023) and we plan to analyze these phenomena in the *Col6a1* Ki Gly292Arg mice.

It is worth mentioning that we did not consider generating a humanized model because the mutation of interest is in an exon and the nucleotide position and corresponding amino acid are highly conserved between human and mice and because of simplicity and cost-effectiveness. Furthermore, considering the high degree of homology between the human and mouse counterparts in this coding region of the *COL6A1* gene, the sequences of nucleic acid-based therapeutics such as antisense oligonucleotides would be highly similar. Thus, the fact that it is not a humanized model does not preclude its value as pre-clinical model.

In summary, the *Col6a1* G292R mice represent the first knock-in model generated by CRISPR/Cas9 for dominant glycine substitutions in the triple helical domain of collagen VI genes which are a common type of mutation in individuals affected with the intermediate and mild forms of COL6-RD. We have quantitatively determined using novel methods several useful molecular, biochemical and functional outcome measures which are relevant to human pathology. This model is therefore a valuable contribution to better understanding the mechanisms of the dominant forms of COL6-RD and to evaluate genetic and non-genetic therapeutic strategies.

## Materials and Methods

### Generation of *Col6a1* knock-in mice using CRISPR/Cas9, genotyping and mouse husbandry

For developing the *Col6a1* (c.874 G>A p.292G>R) mutant mouse model, a *Col6a1*-sgRNA1 5′-CAGGGACGACCTGGGGATCT -3’ targeting the exon 10 was predicted at Breaking-Cas gRNA designer tool (Oliveros, J.C. et al. 2016). The crRNA and tracrRNA (IDT) used to obtain the mature *Col6A1*sgRNA were annealed by mixing equimolar amounts and heating and cooling the mixture to allow hybridization. A designed 200bp ssODN (*Col6a1*ssODN) contained the muted base (c.874 G>A) that mimics the human mutation c.877 G>A (<u>rs398123643:G/R</u>) were produced by chemical synthesis by IDT (Supp.l Table 1). A mixture containing the sgRNA (20 ng/μl), 30 ng/μl of recombinant Cas9 protein (IDT) and 10 ng/μl of the ssODN were microinjected into C57BL6/J zygotes at the Transgenic Facility of the University of Salamanca. Edited founders were identified by PCR amplification (Taq polymerase, NZYtech) with primers flanking the edited region (Suppl Table 1). PCR products were directly sequenced or subcloned into pBlueScript (Stratagene) followed by Sanger sequencing. Selected founders, carrying the desired alleles, were crossed five generations with wild-type C57BL6/6J to eliminate possible unwanted off-targets. Heterozygous mice were re-sequenced and crossed with C57BL6/6J wild-type animals to generate the *BL/6J-Col6a1^em1(c874G>A)Sal^* mouse colony.

For routine genotyping from ear samples, we used the Accustart II Mouse Genotyping Kit, (QuantaBio). DNA was amplified using a pair of primers in the region of the mutation (Supp Table 1) and the 500 bp PCR product digested with Dde1 and the digestion products separated on a 1% agarose gel (see Fig 1). Mice were housed at the registered UB animal facility, where they had ad libitum access to food (regular rodent chow) and water. They were maintained on a light/dark cycle of 08:00–20:00. Both male and female mice were used in equal numbers in the experiments. All experimental procedures involving mice were validated by the local University of Barcelona (UB) Ethics Committees on Animal Experimentation. These procedures were then approved by the Autonomous Government of Catalunya, in accordance with Spanish (RD 53/2013) and European legislation.

### Mouse phenotyping and functional assays

Body weight was measured monthly on the same day. Forelimb maximal strength was assessed using a Grip Strength Meter (IITC Life Science, ALMEMO 2450 AHLBORN, World Precision Instruments, Hertfordshire, UK) with ten measurements per mouse, averaged and normalized to body weight ((https://www.treat-nmd.org/cmd-MDC1A_M.2.2.001). Motor function was tested with a rotarod (Ugo Basile, No. 47600, Comero, Italy), starting at 0 rpm and increasing to 40 rpm. Each test was repeated three times, and the average time on the rotarod was recorded.

### Plethysmography

Respiratory function was assessed using whole-body plethysmography with an EMMS instrument (EMMS, Hants, UK) following standard procedures (https://www.treat-nmd.org/MDX-DMD_M.2.2.002) and the manufacturer’s guidelines. In brief, mice were placed in calibrated chambers containing a pneumatograph that measured pressure differentials within the compartment caused by variations in airflow. The animals were allowed to acclimate in the chambers for 45 minutes at a stable temperature and humidity. Data was collected every 5 seconds using eDacq software (EMMS, Hants, UK). Pause and enhanced pause (Penh) were defined and calculated using the formulas: pause = (Te - RT)/RT and Penh = (PEP/PIP) x pause, where Te is expiratory time, PEP is peak expiratory pressure, and PIP is peak inspiratory pressure. The final values for each parameter were derived from the average of 60 recordings, each of 5 s, representing a total of 5 minutes.

### Tissue collection and processing

Mice (between two and five mice per sex and genotype) were euthanatized by CO2 inhalation followed by cervical dislocation. Dissection and morphological analysis of tissue and muscle samples were performed by veterinary pathologists trained to evaluate mouse tissues. A medial laparotomy and thoracotomy were performed and liver, spleen, left and right kidney, lung, heart and brain were dissected and weighed. Relative weight of each organ was calculated from the body weight (BW) and from the brain weights. Hindlimb and forelimb muscles were dissected and weighed. Muscles were either frozen in dry ice-prechilled isopentane (for histology and immunofluorescence analysis) or snap-frozen in liquid nitrogen (for RNA and protein processing) and stored at -80°C.

### Histology

Hematoxilin and eosin (H&E) staining was conducted on 10um cryosections following standard procedures according to Treat-NMD (https://www.treat-nmd.org/wp-content/uploads/2023/07/cmd-MDC1A_M.1.2.004-68) For the Sirius Red stain sections were incubated in xylene for 10 min and then passed quickly through 100%, 90% and 70% ethanol solutions. Then, sections were incubated in Sirius red solution containing 0.1% (w/v) of Direct Red 80 in picric acid for 60 minutes, washed twice with 2% acetic acid (v/v) and dehydrated serially in graded ethanol, cleared in xylene and mounted in DPX. Histological preparations were imaged on a Nanozoomer S60 (Hamamatsu).

### RNA isolation and Reverse Transcription

Extraction and purification of total RNA from cell cultures were performed with the RNeasy® Fibrous Tissue Mini Extraction Kit (Qiagen, Hilden, Germany). RNA concentration was measured as for DNA. Samples were stored at 80 °C. For reverse transcription, equal amounts of RNA (300ng) were used and added to an M-MLV reverse transcriptase reaction mix (Promega, Madison, WI, USA).

### Digital droplet PCR

Digital PCR was performed as previously described (López-Márquez et al., 2022) in a reaction volume of 20uL using SuperMix for Probes (no dUTP) (Bio-Rad, Hercules, CA, USA), 450 nM of each primer pair, 250 nM of each probe (Suppl. Table 1) and 0.025 or 2 ng of cDNA depending on whether allele-specific or total expression is measured. PCR cycling conditions were optimized for each primer pair. Data were analyzed using Bio-Rad QuantaSoftTM software (v1.7.4), (Bio-Rad, Hercules, CA, USA) with default settings for threshold determination to distinguish positive and negative droplets. The wild type and mutant alleles were distinguished by a 2-dimensional view of the ddPCR analysis. At least three biological replicates were carried out for each experiment.

### Protein isolation, quantification and analysis

Approximately twenty to sixty mg of muscle tissue was mechanically homogenized using a Tissue Ruptor (Qiagen) in lysis buffer containing 75mM Tris-HCl pH 6.8, SDS10%, NAF (10mM), Na3VO4 (1mM), and a cocktail of protease inhibitors (Roche, ref-04693159001). Then we centrifuged the lysates at 4°C and 14000 g for 15 min. Protein concentration in the lysates was measured using the Pierce BCA Protein Assay (Thermo Scientific, ref. 23223 and 23224). Protein lysates (40ug per lane) were separated under reducing conditions on a Bis-Tris 4–12% polyacrylamide precast gel (Bio-Rad, Hercules, CA, USA). Blotted membranes were blocked for 1 h in 5% milk or BSA in Tris-buffered saline (10 mM Tris, 150mMNaCl, pH 8.0) plus 0.1% Tween, and incubated overnight at 4°C with rabbit anti-collagen VI (70R-CR009x, Fitzgerald) or rabbit anti-desmin (ab8592, Abcam). Membranes were washed and incubated for 1 h at room temperature with peroxidase-conjugated secondary antibodies (1:10.000, Jackson ImmunoResearch Laboratories, USA). Bands were revealed with the ECL chemiluminescence detection system (Thermo Scientific, Waltham, MA, USA) using the iBright detection equipment (Invitrogen) and the density of bands quantified using Fiji ImageJ software.

### Immunofluorescence

Cryosections (10 um) were fixed in 4% paraformaldehyde for 10 min at room temperature and washed twice in PBS-0.1M glycine solution, followed by two washes in PBS alone before being blocked for 30 min in 5% normal goat serum solution containing 0.5% triton in PBS for 60 min. A mixture of primary antibodies [rabbit anti-collagen VI 1:1000 (ab6588, Abcam) and rat-anti-perlecan 1:5000 (MAB1948, Merck)] in the blocking solution was added to the sections overnight at 4°C. Sections were washed in PBS and incubated in a mixture of secondary antibodies (Alexa Fluor® 488 goat anti-rabbit IgG 1:500 and Alexa Fluor® 594 goat anti-rat IgG 1:500) in the blocking solution containing Dapi for 30min at room temperature. Finally, sections were washed three times with PBS and mounted with Fluoromount-G®.

### Automated Muscle Tissue Mapping with Confocal Microscopy

Confocal microscopy analysis was conducted using a Leica TCS SP8 STED 3X system, equipped with a white light laser, HyVolution mode, and hybrid detectors (Leica Microsystems, Wetzlar, Germany). A high-precision motorized stage, controlled by LASX Navigator software, was employed to capture large-scale 3D mosaics of tissue sections at low magnification (HC PL APO CS2 10×/0.4 dry objective). The software automatically calculated the optimal stage positions based on the image dimensions in microns and the degree of overlap between adjacent images, ensuring complete coverage of the volume of interest. Individual image tiles were captured at 1024 × 1024 pixels with a z-step of 5 μm and 12 bits, and 4–20 stacks were collected for each extended image. Muscle tissues were sequentially excited at three wavelengths: 405 nm (DAPI), 488 nm (COL6), and 594 nm (Perlecan). Detection ranges were 420–465 nm for DAPI, 500–560 nm for COL6, and 610–755 nm for Perlecan. Ten sections were acquired every 5 μm across the tissue thickness, and maximum projections were generated using LAS AF™ software (Leica Microsystems, Heidelberg, Germany).

### Image Analysis

For the analysis of muscle fiber morphology, nuclei, collagen VI, and perlecan intensity, images were extracted using Leica Application Suite X (LAS X) and processed through the Cellpose (Stringer et al., 2021) model, which uses α convolutional neural network to automatically deblur, denoise, and segment fibers. Post-processing steps removed irregularly shaped fibers, those with non-smooth borders, and isolated fibers to ensure reliable segmentation. The post-processed segmentations were saved as masks and divided into patches to enhance data availability. For muscle fiber size analysis and internal nuclei quantitation, the original images and masks were processed to calculate the minimum Feret diameter and internal nuclear density per fiber for each fiber within the patches. For intensity analysis, the original images and masks were imported into QuPath (Bankhead et al., 2017), where fluorescence intensity was measured at the fiber periphery (basal lamina) within a 2 µm area. Results were exported to an Excel file, and scripts are available upon request.

### Quantification of collagen area/fibrosis

To quantify the fibrotic tissue area from the muscle sections an automatic image analysis pipeline has been programmed with FIJI-ImageJ Software. Given the irregularity of some muscle sections, the analysis was performed by selecting specific Regions of Interest (ROI) of 500x500 pixels. This process was done by manually moving a rectangular window over the muscle section. When the window was placed on top of a desired ROI, the algorithm automatically segments the fibrotic area of the muscle using the Li algorithm. Then, the area of the segmented region and its percentage over the whole ROI were computed. This process can be repeated by manually moving the window to select all the muscle ROIs that must be quantified to map the whole muscle (Schindelin, J. et al., 2019). Between 41 and 176 ROIs were analysed depending on the size of the muscle section.

### Statistical Analysis

For the analysis of the body weight, piecewise regression was used to study the age at which the weight growth changes, after visually detecting this tendency with the corresponding boxplots. After the determination of the age range for these two time periods with clear different weight evolutions, they were analysed separately. In both cases, linear regression models were used to study how age, sex and genotype explained weight for all the data. This analysis was also performed for both sexes separately. Plethysmography data was analysed with linear mixed models, in which age, sex and genotype were included as fixed effects and random effects associated to subject ID were defined. Normality and homocedasticity in the residuals from these models were assessed, using robust methods to create linear mixed models in cases where the residuals showed problems in the model fitting. All p-values under 0.05 were considered significant. Bonferroni-Holm correction for multiple testing was applied to the results from the plethysmography models. For the morphometric analysis of muscle fibres (Minimum Feret diameter and number of internal nuclei) and the analysis of intensity of collagen VI and perlecan immunofluorescence numerical variables, data was described with the appropriate descriptive statistics. For each variable, different genotype groups were compared with Kruskal-Wallis test, using pairwise Mann-Whitney’s U-test with Bonferroni-Holm adjustment as post-hoc analysis. r parameter was used as effect size measure for these comparisons. Significant results with effect sizes over 0.1 were considered relevant and discussed. Differences between groups in Minimum Feret diameter distribution were studied comparing 95% confidence intervals of the variance coefficient of each group.

One-way ANOVA test was used followed by Tukey’s HSD as a post-hoc test to determine statistical significance for the comparison of the weight of the isolated muscles, the grip strengths test, the digital PCR, Western blot and percentage (%) area of fibrosis values. Statistical analyses and graphs were performed using R (v. 4.3), working with RStudio (v. 2022.2) or GraphPad PRISM Version 10.4.0.

## Supporting information

Supplementary Image 1

## Acknowledgements

1. P. H-C. and M. S-M. would like to thank Ignacio García-Tuñon for his assistance in CRISPR technology.

## Data and resource availability

All original data are available from the authors without any restrictions.

## Author Contributions

Conceptualization: CJ-M, AL-M, M S-M, SB, MR.

Formal analysis: CJ-M, A L-M, BC, ZG, VA, SB, AQ, MR.

Investigation: CB, LLE, P H-C, M S-M, IG, MB-R, EP, MR, A L-M, C J-M.

Writing – original draft: C J-M, A L-M.

Writing-review and editing: all authors. Visualization: C J-M, A L-M, SB, MR.

Supervision: C J-M, A L-M.

Funding acquisition: C J-M, MS-M, MR.

---

Supplementary Fig 1: Hematoxilin and eosin (H&E) staining of 12 months old mice. Scale bar= 100µm.

Supplementary Fig 2. Box-plots of collagen VI/perlecan intensity ratio (median and interquartile range). A.u. = arbitrary units.

Supplementary Table 1: The sequence of the primers used to genotype mice by PCR as well as the sequence of the primers and probes (FAM or HEX labelled) used to amplify and detect the wild-type and mutant alleles by ddPCR is indicated. For gene expression analysis of col6a transcripts we used Gene Expression Assays by Thermofisher Scientific (Inventoried Best Coverage) that contain the forward and reverse primers and the corresponding probes (FAM labelled). We also indicate the sequence of the single stranded DNA template (ssODN) that was used to generate the col6a1 Ki mice.

Supplementary Table 2: Summarizes the statistical analysis of the body weight. We used linear regression. Age, sex and genotype were included as independent variables. All p-values under 0.05 after Bonferroni-Holm correction for multiple testing were considered significant.

Supplementary Table 3: Summarizes the statistical analysis of the plethysmography data. Linear mixed models were used. Age, sex and genotype were included as independent variables in these models, and subject ID were used to define random effects. All p-values under 0.05 after Bonferroni-Holm correction for multiple testing were considered significant.

Supplementary Table 4: Coefficient of Variation for the minimum Feret‗s diameter (mean and 95% confidence interval) in 6 months old mice. Only muscles where there was no overlap in the CI between genotypes were considered relevant.

Suppl. Table 5. Analysis of the collagenVI/perlecanfluorescence intensity ratio between genotypes in 12 months old mice. A. Mean intensity in A.u. (and standard deviation) are indicated for each genotype. We used Kruskal-Wallis test to compare the values from these 3 groups, using Mann-Whitney’s U-test with Bonferroni-Holm correction for multiple testing as post-hoc analysis. The p value is indicated for the comparison between Wild-type (wt) and heterozygous (het) in the het column and for the comparison between wild-type and homozygous (hom) in the hom column and in brackets the value for the comparison het versus hom. We also calculate the r parameter as an effect size measure for these tests, as we have a large sample size, and it is relatively easy to obtain significant results with really small differences between groups. * An r (effect sizes) equal or larger than 0.1i s considered relevant. The r parameter is indicated for the comparison between wt and het in the het column and for the comparison between wt and hom in the hom column and in brackets the value for the comparison het vs hom. Only r values = or > than 0.1 are indicated. Fold change between wt and the other genotypes is also indicated (wt/het or hom).

