## Supplementary Table 1 for "Generation and Characterization of *Col6a1* knock-in mice: A Promising Pre-Clinical Model for Collagen VI-Related Dystrophies"

**Suppl Table 1**.

| Gene/allele | Primer | Sequence (5 ´to 3´) | Probe/allele | Sequence (5 ´to 3´) |
|---|---|---|---|---|
| Col6a1 genotyping (Salamanca) | Forward | CTCCCTTAAGGATTCTGTGTGG | | |
| Col6a1 genotyping (Salamanca) | Reverse | GAGAAGGCAAGGAGGGTCTTC | | |
| Col6a1 genotyping (Barcelona) | Forward | GGTGCATAACCTCTCCTCCA | | |
| Col6a1 genotyping (Barcelona) | Reverse | AGAAGGCAAGGAGGGTCTTC | | |
| col6a1 wild type/mutant | Forward | CGAGGAAAGCCAGGTCTTC | Wild type | ATCTTGGACCAGT (HEX) |
| col6a1 wild type/mutant | Reverse | TTCTCCCTTCATACCCTGGTA | Mutant | ATCTTAGACCAGTC (FAM) |
| **Gene** | **Gene Expression Assay (ref)** | **Sequence (5 ´to 3´)** | | |
| Col6a1 | Mm_00487160_m1 | N/A | | |
| Col6a2 | Mm_00521578_m1 | N/A | | |
| Col6a3 | Mm_00711678_m1 | N/A | | |
| Col6a5 | Mm_01231908_m1 | N/A | | |
| Col6a6 | Mm_00556810_m1 | N/A | | |
| **Gene/allele** | **ssODN** | **Sequence (5´to 3´)** | | |
| **col6a1** | | CTGTAGCAAGGGGTAAGGTGGGCTGAGAAGGACCTTCCAGTGCTATCCAGGGATGTCT GACTATCTCTTGTTGTGTTCCAGGGACGACCTGGCGATCTTAGACCAGTCGGGTATCAG GGTATGAAGGTACGTGTCCTTAATGTCCAATGGCCTCTCTTGGGTGGGGTGGATCAGGC TTTGGAAGCAAAGCCCAAGCTGCC | | |
