## Supplementary Table 4 for "Generation and Characterization of *Col6a1* knock-in mice: A Promising Pre-Clinical Model for Collagen VI-Related Dystrophies"

|  | **CV (95% CI interval)** | | |
| --- | --- | --- | --- |
|  | **Wt** | **Het** | **Hom** |
| *M Gastrocnemius* | 0.338(0.333-0.343) | 0.357 (0.352-0.362) | 0.331 (0.324-0.339) |
| *M Tibialis anterior* | 0.249(0.243-0.255) | 0.334(0.328-0.34) | 0.384(0.365-0.405) |
| *F Triceps* | 0.247(0.242-0.253) | 0.265(0.26-0.27) | 0.305(0.293-0.317) |

*Suppl. Table 4.* ***A****. Coefficient of Variation for the minimum Feret´s diameter (mean and 95% confidence interval) in 6 months old mice.*
