## Supplementary figures and images for "Generation and Characterization of *Col6a1* knock-in mice: A Promising Pre-Clinical Model for Collagen VI-Related Dystrophies"

### Supplementary Fig 2

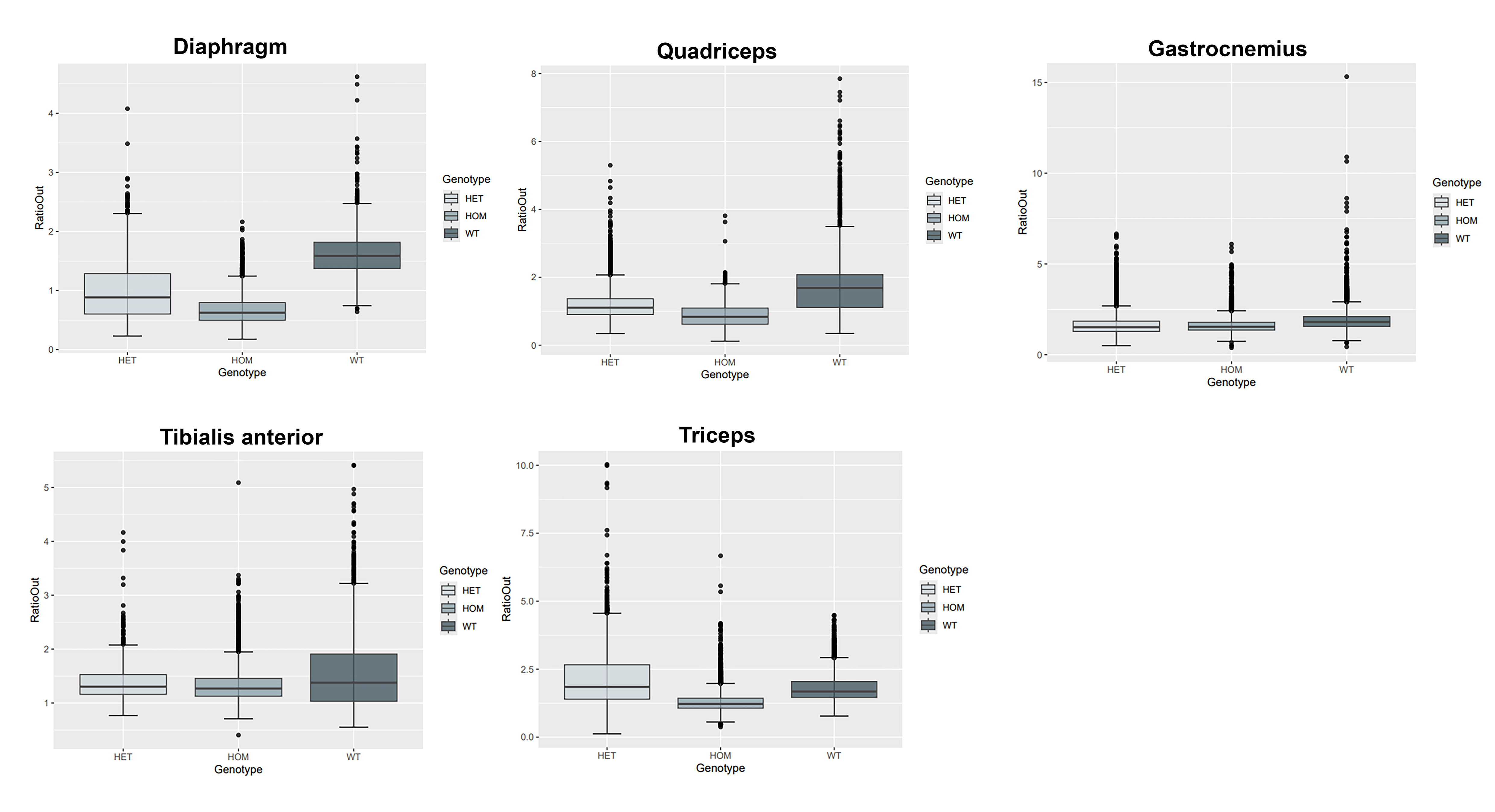

### Supplementary Image 1

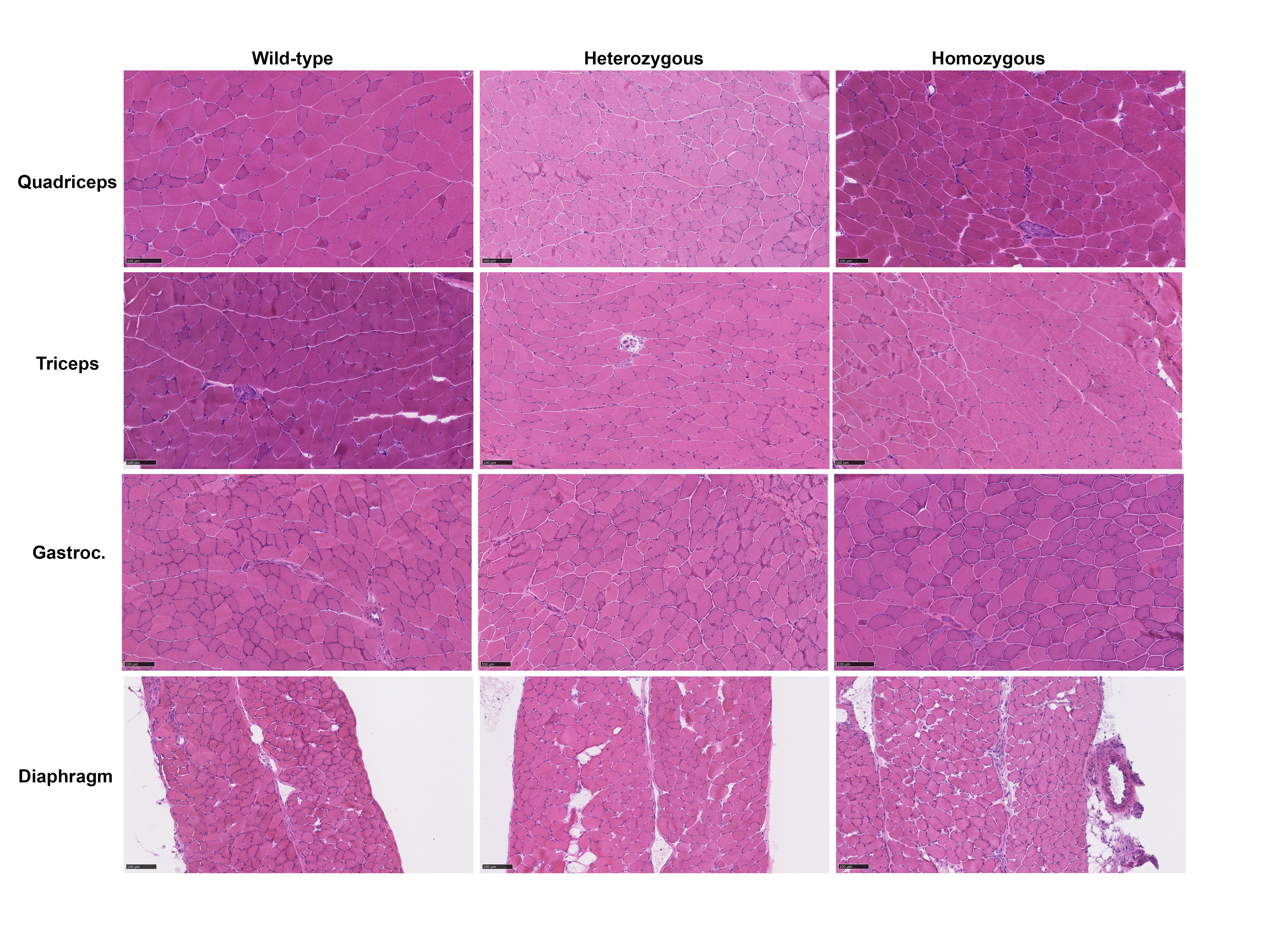

### Supplementary Table 5

Suppl Table 5

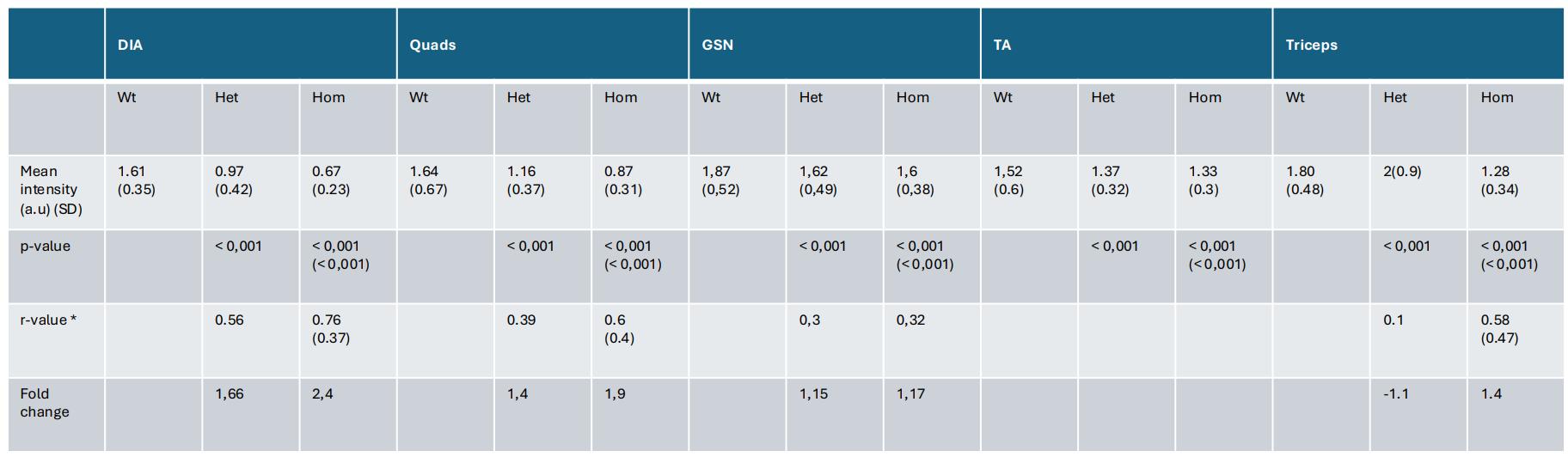
